# RETINOBLASTOMA RELATED (RBR) interaction with key factors of the RNA-directed DNA methylation (RdDM) pathway

**DOI:** 10.1101/2022.01.06.474281

**Authors:** León-Ruiz Jesús, Espinal-Centeno Annie, Blilou Ikram, Scheres Ben, Arteaga-Vázquez Mario, Cruz-Ramírez Alfredo

## Abstract

- Transposable elements and other repetitive elements are silenced by the RNA-directed DNA methylation pathway (RdDM). In RdDM, POLIV-derived transcripts are converted into double stranded RNA (dsRNA) by the activity of RDR2 and subsequently processed into 24 nucleotide short interfering RNAs (24 -nt siRNAs) by DCL3. 24-nt siRNAs are recruited by AGO4 and serve as guides to direct AGO4 - siRNA complexes to chromatin bound POLV-derived transcripts generated from the template/target DNA. The interaction between POLV, AGO4, DMS3, DRD1, RDM1 and DRM2 promotes DRM2-mediated *de novo* DNA methylation.
- The Arabidopsis Retinoblastoma protein homolog is a master regulator of cell cycle, stem cell maintenance and development. *In silico* exploration of RBR protein partners revealed that several members of the RdDM pathway contain a motif that confers high affinity binding to RBR, including the largest subunits of POLIV and POLV (NRPD1 and NRPE1), the shared second largest subunit of POLIV and POLV (NRPD/E2), RDR1, RDR2, DCL3, DRM2 and SUVR2. We demonstrate that RBR binds to DRM2, DRD1 and SUVR2. We also report that seedlings from loss -of-function mutants in RdDM and in *RBR* show similar phenotypes in the root apical meristem. Furthermore, we show that RdDM and SUVR2 targets are up-regulated in the *35S::AmiGO-RBR* background.
- Our results suggest a novel mechanism for RBR function in transcriptional gene silencing based on the interaction with key players of the RdDM pathway and opens several new hypotheses, including the convergence of RBR-DRM2 on the transcriptional control of TEs and several cell/tissue and stage -specific target genes.

## Introduction

DNA methylation is essential for proper development in eukaryotes. In plants, it is involved in the regulation of gene expression, the defense against invasive nucleic acids, both of them with effects on development and physiology. In plants, cytosines can be methylated in symmetrical (CG or CHG) and asymmetrical (CHH) sequence contexts (where H can be A, T, or C). Transposable elements (TEs) and other repetitive sequences are the main targets of DNA methylation (Borges and Martienssen, 2015; Matzke and Mosher, 2014). The major small RNA-mediated epigenetic pathway involved in *de novo* DNA methylation is the RNA-directed DNA methylation (RdDM) pathway (Matzke and Mosher, 2014; Erdmann and Picard, 2020). RdDM involves the function of Nuclear RNA Polymerase D (NRPD) or POL IV and NRPE or POLV (Hagg and Pikaard, 2011). POLIV transcribes short single stranded RNA (ssRNA) 26 to 45 nt in length (from the target locus that will be methylated) that serve as substrate for RNA-DEPENDENT RNA POLYMERASE 2 (RDR2) for the generation of double stranded RNA (dsRNA). The resulting dsRNA is processed by DICER-LIKE 3 (DCL3) into 24-nt small interfering RNAs (siRNAs). HUA ENHANCER 1 (HEN1) methylates 24-nt siRNAs at their 3’-end and are subsequently recruited by ARGONAUTE 4 (AGO4) (or other close paralog such as AGO6 and AGO9). The AGO4-siRNA complex associates with chromatin bound POLV-dependent transcripts produced from the same loci that will be methylated, through RNA-RNA pairing. The association between the AGO4-siRNA complex and POLV is further stabilized by protein-protein interactions between AGO4 and the CTD of POLV. Recruitment of the *de novo* DNA methyltransferase DOMAINS REARRANGED 2 (DRM2) to the template/target DNA occurs through the activity of RNA-DIRECTED DNA METHYLATION 1 (RDM1) that is able to bind methylated single stranded DNA (ssDNA) and also interacts with DRM2 and AGO4 (reviewed in Matzke & Mosher, 2014; Trujillo et al., 2018).

Retinoblastoma proteins are multi-faceted master regulators of cell reprogramming in eukaryotes and are involved in the control of cell cycle, DNA damage response and in protein-protein interactions (PPIs) with transcription factors that modulate stem cell maintenance and asymmetric cell division for proper cell lineage commitment (Calo et al., 2010; Cruz-Ramirez et al., 2012; Harashima & Sugimoto, 2016; reviewed in Dyson, 2019; reviewed in Desvoyes & Gutiérrez, 2020). In Arabidopsis, RETINOBLASTOMA RELATED (RBR) has been shown to bind DNA, putatively to regulate the transcription of hundreds of genes and transposable elements (Bouyer et al., 2018) and also indirectly modulates gene expression by PPIs and genetic interactions with lineage-specific transcription factors (Cruz-Ramirez et al., 2012; Cruz-Ramirez et al., 2013; Matos et al., 2014; Zhao et al., 2017), chromatin-remodeling factors such as PICKLE (PKL) (Ötvös et al., 2021), and the Polycomb Repressor Complex 2 (PRC2) (Julien et al., 2018). The PRC2 complex regulates plant growth and development through the trimethylation of Lysine 27 on Histone 3 (H3K27me3), a well-known epigenetic mark involved in transcriptional repression. Two independent studies have established the connection between RBR and PRC2. Jullien *et al*., (2008) demonstrated that RBR directly binds to MULTICOPY SUPPRESSOR OF IRA1 (MSI1), an essential component of *Arabidopsis* PRC2 protein complexes involved in female gametogenesis, seed and vegetative development. The RBR-MSI1 complex directly represses *DNA METHYLTRANSFERASE 1* (*MET1*) transcription, MET1 is a DNA methyltransferase acting on cytosine methylation at symmetrical CpG positions. *MET1* repression occurs only on the female gamete and is required for the expression of imprinted genes. A similar observation was also reported by Johnston *et al*. (2008). The interaction between RBR and PRC2 is potentially deeper since FERTILIZATION-INDEPENDENT ENDOSPERM (FIE), another member of the PRC2 complex that interacts with MEDEA (MEA), SWINGER (SWN) and CURLY LEAF (CLF) (Oliva *et al*., 2016) does contain a highly conserved LxCxE motif, which is characteristic of proteins that bind with high-affinity to RBR (Cruz-Ramírez, *et al*., 2012).

Plant and animal Retinoblastoma proteins share conserved residues that allow them to interact with proteins containing an LxCxE SLiM (SLiM: Short Linear Motif) RBR-binding motif (*Lee et al*., 1998, Dick, 2007). A decade ago, a global search in the *Arabidopsis* proteome for proteins containing the LxCxE SLiM led us to the identification of hundreds of candidates that potentially interact with the single *Arabidopsis* Retinoblastoma protein: RBR. By employing the LxCxE motif, that confers high-affinity to RBR, as an *in silico* bait to identify *Arabidopsis* RBR protein partners (Cruz-Ramirez *et al*., 2012), we identified several components of the RdDM pathway including the largest subunits of POLIV and POLV, RDR1, RDR2, DCL3, DRM2 and SUVR2 as potential targets of RBR. In this study we demonstrate that RBR binds to DRM2, DRD1 and SUVR2. We also report that seedlings of loss-of-function mutants in RBR and in genes of the RdDM pathway show phenotypes in the root apical meristem, with defects in the RSCN. This is consistent with the observation that RdDM and SUVR2 targets are up-regulated when RBR is post-transcriptionally silenced using the cell-type-specific artificial microRNA for Gene-silencing Overcome (amiGO) system. Our results uncover a novel mechanism for RBR function in transcriptional silencing through its interactions with key components of the RdDM pathway and opens the possibility of a convergent action of RBR-DRM2 in the regulation of TEs and lineage or tissue-specific transcription factors, and stem cell regulators, such as *WUSCHEL, AGL15 and POLAR*, among other interesting putative target genes.

## Materials and Methods

### Plant Materials

*Arabidopsis thaliana* plants were grown as described in Cruz-Ramirez *et al*. (2004). Col-0 wild type, double (*nrpd2a-2;nrpd2b-1)* and triple mutants (*drm1;drm2;cmt3*) plants were used for phenotypic analyses, as well as transgenic lines (*pRBR::RBR:CFP, pDRM2::DRM2-GFP* and *35S::AmiGORBR)* (Cruz-Ramirez *et al*., 2012; Cruz-Ramirez *et al*., 2013).

### Microscopic Analysis

Seedlings were grown and roots were prepared for confocal microscopy as previously described (Cruz-Ramirez *et al*., 2012). Fluorescent signals for the diverse genetic backgrounds were recorded with a Leica SP2 CLSM and a Zeiss LSM 800 CLSM. Roots were mounted and stained with Lugol as in Willemsen *et al*. (1998) and were visualized by Nomarski optics.

### Protein-Protein interaction (PPI) assays

Yeast two-hybrid (Y2H) interactions were characterized by employing the ProQuest Two-Hybrid System (Invitrogen Life Technologies) as reported in Cruz-Ramirez *et al*., (2013). To quantify the strength of each interaction, three biological and technical replicates of Beta-galactosidase assays with CPRG as substrate were performed. Bimolecular Fluorescence Complementation Assays in *Arabidopsis* protoplast were performed as reported in Cruz-Ramirez *et al*., (2012). For RBR-DRM2, RBR-DRD1 and controls YFP fluorescence was recorded with a Leica SP2 CLSM.

### Computational analyses and ortholog identification

Angiosperm protein sequences were downloaded from Phytozome (https://phytozome-next.jgi.doe.gov/), while non-angiosperm and algae protein sequences were downloaded from Phytozome, Fernbase (Li *et al*., 2018), TreeGenes (Wegrzyn *et al*., 2019) and Phycocosm (Grigoriev *et al*., 2021). Sequences for *A. agrestis* and *P. margaritaceum* were downloaded directly from the University of Zurich Hornworts database (Li *et al*., 2020: https://www.hornworts.uzh.ch/en.html) and the Penium genome database (Jiao *et al*., 2020: http://bioinfo.bti.cornell.edu/cgi-bin/Penium/blast.cgi), respectively. LxCxE-SLiM containing protein sequences were detected using a custom perl script (Caballero-Perez, personal communication). To infer orthologues, all protein sequences from all 28 species analyzed were placed into orthogroups using the OrthoFinder software (Emms *et al*., 2019).

### qRT-PCR assays of RdDM targets

Twenty seedlings of 14-days-old post germination plants from Col-0 or *35S::AmiGORBR* (Cruz-Ramírez *et al*., 2013), were used for total RNA extraction by TRIzol reagent (ThermoFisher) in three biological replicates. Total RNA was used to generate cDNAs according to the manufacturer’s protocol for SuperScript lll (ThermoFisher) we used 5 μg of total RNA per 20 μL reaction. The expression level was determined using SYBR GREEN mix (ThermoFisher) in a 10 μL reaction. The data were normalized using Actin 7 expression levels. The primers used in these experiments are those reported in Han *et al*. (2014).

## Results and discussion

### Major players of the RdDM pathway and their putative RBR-Binding motifs

Early predictions for Arabidopsis RBR-interactors, served as the basis for the functional characterization of the interaction between RBR with diverse lineage-specific factors such as SCARECROW, FAMA, XND1 and PICKLE, among others (Cruz-Ramirez *et al*., 2012; Matos *et al*., 2014; Zhao *et al*., 2017; Zhou *et al*., 2019; Ötvös, *et al*., 2021). In addition to the aforementioned proteins, we identified many proteins with diverse key molecular and cellular functions bearing the RBR-binding motif which, in many cases, were evolutionarily conserved. Among them, we found that components of the RdDM pathway including NRPD2, DRD1, DRM2, DCL3 and SUVR2 contain the canonical LxCxE SLiM (Fig. 1, TableS1). We also found that major players of the RdDM pathway including NRPD1, NRPE and RDR2 contain a non-canonical RBR-interaction motif I/LxFxE (Fig. 1, TableS1). The observation that eight components of the RdDM pathway shared canonical and non-canonical RBR-interaction motifs prompted us to investigate if some of them are true physical RBR interactors.

**Fig. 1.**
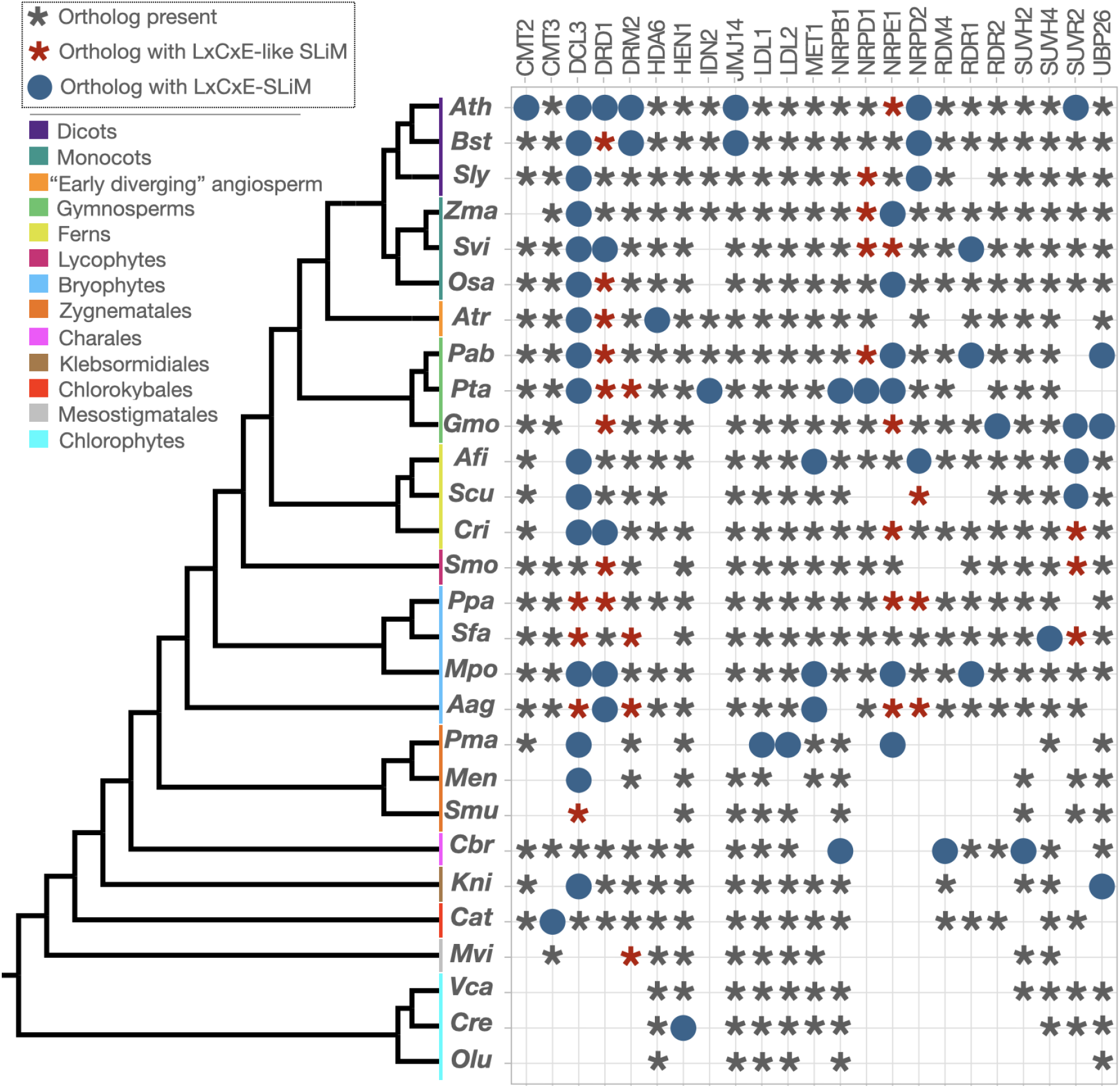
Phylogenetic conservation of canonical (solid blue circle) and non-canonical (red asterisk) LxCxE SLiMs in orthologs of the RdDM pathway in representative species along Viridiplantae.

### Conservation of LxCxE-like motifs in RdDM factors along Viridiplantae

To gain insight into the evolutionary conservation of the LxCxE SLiM present in components of the RdDM pathway, we interrogated publicly available plant and algae genomes aiming to detect the presence of canonical and non-canonical (IxCxE/LxCxD/IxCxD) LxCxE SLiMs among orthologs of the RdDM pathway along the Viridiplantae clade. To optimize the breadth of the plant phylogeny to cover, we focused on a small subset of species with available sequenced genomes representing each major lineage of the Viridiplantae kingdom. The species selected and analyzed include representatives from angiosperms (*A. thaliana* [*Ath*]; *Boechera stricta* [*Bst*]; *Solanum lycopersicum* [*Sly*]; *Zea mays* [*Zma*]; *Setaria viridis* [*Svi*]; *Oryza sativa* [*Osa*]; *Amborella trichopoda* [*Atr*]), gymnosperms (*Picea abies* [*Pab*]; *Pinus taeda* [*Pta*]; *Gnetum montanum* [*Gma*]), ferns (*Azolla filliculoides* [*Afi*]; *Salvinia cucullata* [*Scu*]; *Ceratopteris richardii* [*Cri*]), lycophytes (*Selaginella moellendorffii* [*Smo*]), bryophytes (*Sphagnum fallax* [*Sfa*]; *Physcomitrium patens* [*Ppa*]; *Marchantia polymorpha* [*Mpo*]; *Anthoceros agrestis* [*Aag*]) charophyte (*Penium margaritaceum* [*Pma*]; *Mesotaenium endlicheranium* [*Men*]; *Spirogloea muscicola* [*Smu*]; *Chara braunii* [*Cbr*]; *Klebsormidium nitens* [*Kni*]; *Chlorokybus atmophyticus* [*Cat*]; *Mesostigma viride* [*Mvi*]) and chlorophyte (*Volvox carteri* (*Vca*); *Chlamydomonas reinhardtii* [Cre]; *Ostreococcus lucimarinus* [*Olu*]) algae.

Our analysis revealed that *A. thaliana* was the species with more proteins containing either canonical or non-canonical RBR-binding motifs (Fig. 1, Table S1), with 8 out of 23 RdDM-related proteins analysed (DCL3, DRD1, DRM2, NRPD2, SUVR2, CMT2, JMJ14 and NRPE1). DCL3 orthologs showed the highest level of conservation for canonical and non-canonical LxCxE SLiMs among the species analyzed as they are absolutely conserved in tracheophytes, with the only exception of *G. montanum*. Interestingly, while DCL3 in *M. polymorpha* bears a canonical LxCxE SLiM, DCL3 orthologs in other bryophytes specifically *S. fallax, P. patens* and *A. agrestis* contain LxCxE-like SLiMs. From the seven charophyte algae species analysed, 3 of them contain canonical RBR-binding motif (*P. margaritaceum M. endlicheranium, K nitens*) while *S. muscicola* contains an LxCxE SLiM (Fig. 1, Table S1). Although the LxCxE SLiM is highly conserved along DCL3 orthologs, it is difficult to determine if the canonical or the non-canonical motif is the ancestral one.

DRD1 orthologues showed the presence of the LxCxE SLiM in a patchy pattern along the plant lineages analyzed. The presence of the LxCxE SLiM in DRD1 orthologues is less conserved than in DCL3 orthologues since we were not able to find LxCxE or LxCxE-like SLiMs in any of the algae species analyzed, however it is present in *Marchantia, Anthoceros*, and *Ceratopteris* DRD1 orthologs (Fig. 1, TableS1). The presence of the LxCxE SLiM is even less conserved in DRM2 orthologs than in DRD1, with only two DRM2 orthologs from *Arabidopsis* and *Boechera* exhibiting a canonical SLiM and non-canonical LxCxE SLiMs present in *Pinus, Sphagnum, Antoceros* and *Mesostigma*. In the case of the subunits of POLIV and POLV, we expanded a presence-absence analysis along the plant phylogeny, similar to that reported previously by Huang *et al*. (2015). We found that NRPE1, NRPD1 and NRPD2 showed the presence of both canonical and non-canonical LxCxE SLiMs in diverse species, among these 3 proteins we found that NRPE1 is the one with more species containing either canonical or non-canonical RBR-binding SLiM (Fig. 1, TableS1). While Arabidopsis NRPD1 does not contain an LxCxE SLiM, *P. taeda* NRPD1 ortholog contains a canonical LxCxE SLiM and orthologs from *S. viridis, P. abies*, maize and tomato bear a non-canonical LxCxE SLiM. We observed the presence of canonical LxCxE SLiMs in NRPE1 from charophyte to flowering plants (*P. margaritaceum, M. polymorpha, P. abies, P. taeda, Z. mays and O. sativa)* and non-canonical LxCxE SLiMs in NRPE1 orthologs from *C. richardii, A. agrestis, P. patens, G. montanum, S. viridis and A. thaliana*. The presence of canonical and noncanonical LxCxE SLiMs involved in RBR-binding in the POLIV and POLV largest subunits (NRPD1 and NRPE1, respectively) and the shared second largest subunit (NRPD/E2) strongly suggests that a new layer of regulation of the RdDM pathway mediated by RBR is present in land plants.

### DRM2, DRD1 and SUVR2 physically interact with RBR

Based on their conservation patterns we selected a group of proteins to test for protein-protein interactions with RBR. We generated constructs using amplified coding sequences (CDS) of DRM2, DRD1 and SUVR2 from *Arabidopsis* for Y2H assays, in order to test if they interact with RBR (previously cloned in pDEST32 and used in Cruz-Ramirez *et al*., 2012). Our results showed that SUVR2 strongly interacts with RBR when quantified and compared with other partners and controls (Fig.2 a, b) but DRD1 and DRM2 showed weak interaction. The previously described Y2H results prompted us to confirm, using a semi *in vivo* system, DRD1-RBR and DRM2-RBR interactions by Bimolecular Fluorescence Complementation (BiFC) assays. We found that YFP nuclear signal is clear and evident in *Arabidopsis* mesophyll protoplasts, confirming that DRD1 and DRM2 do interact with RBR. We also found that the *M. polymorpha* DCL3 ortholog interacts with both Arabidopsis and *M. polymorpha* RBRs by Y2H assays (León-Ruiz & Cruz-Ramirez, *in preparation*). Further experimental work is required to confirm PPIs between RBR and other RdDM-related proteins including NRPD1, NRPE1 and NRPD/E2 but it is important to consider that regulation by RBR can go beyond its direct interactors, for example it can affect other PPIs indirectly as documented in the IntAct Molecular Interactions Database from EMBL-EBI: DRM2 establishes 12 PPIs, from which at least 4 are direct interactions with members of the RdDM pathway, such as RDM1, AGO4, AGO9, and ZOP1 (Fig.4d. (https://www.ebi.ac.uk/intact/interactions?conversationContext=4). Taken together, our results indicate that the evolutionary conservation of LxCxE SLiMs among components of the RdDM pathway is consistent with our experimentally validated interactions with RBR in the cases of DRD1, DRM2 and SUVR2.

**Fig. 2.**
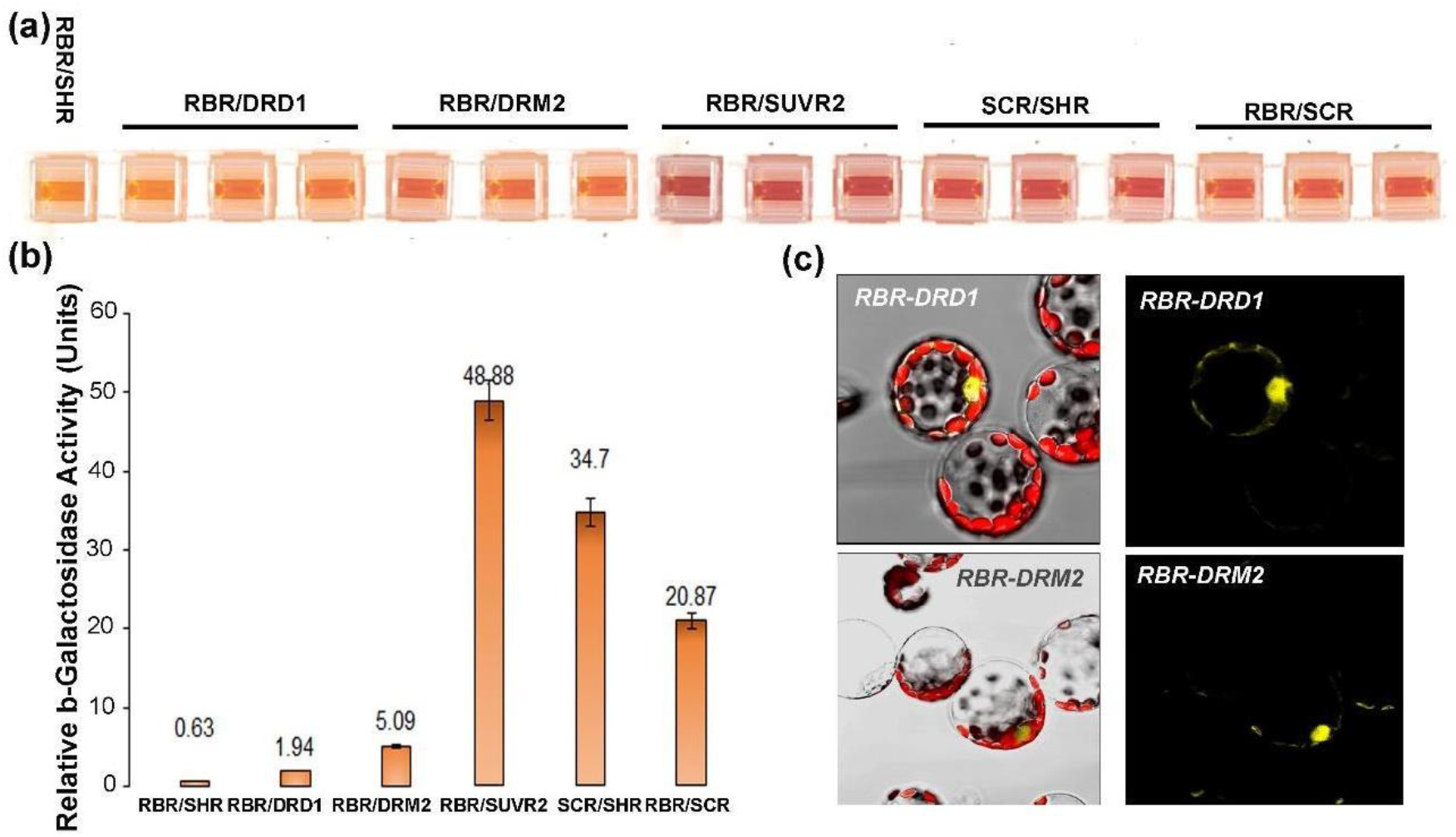
(a) Yeast two-hybrid analyses showing b-gal colorimetric reaction and its quantitation, in (b), for diverse proteins of the RdDM pathway and RBR. SCR-RBR and SCR-SHR combinations are positive controls, and RBR-SHR is the negative control (Cruz-Ramírez, etal, 2013) (c) RBR-DRD1 and RBR-DRM2 binding by BiFC in *Arabidopsis* mesophyll protoplasts.

### RdDM and RBR loss-of-function mutants show similar developmental alterations

It has been shown that loss of function mutants in members of the RdDM pathway show phenotypes in diverse developmental processes and stages of *Arabidopsis* (He *et al*., 2009; reviewed in Matzke *et al*., 2015; Mendes *et al*., 2020).

In addition to physically interacting, RBR and DRM2 protein fusions (pRBR::RBR:CFP, pDRM2::DRM2:GFP) have quite similar expression patterns as both proteins are present in every cell of the RAM (Fig.3 a,b). Since RBR has been shown to regulate stem cells and QC divisions in the Arabidopsis RAM (Cruz - Ramirez *et al*., 2012; Cruz-Ramirez, *et al*., 2013), we wondered if loss of function (LOF) mutants, in tested and putative interactors, in genes of the RdDM pathways may display similar phenotypes to those in RBR LOF lines. Therefore, we analyzed root development of 12 dpg (days post germination) seedlings of the *drm1;drm2;cmt3* triple mutant and the *nrpd2a;nrpd2b* double mutant and observed that primary root development in these mutants is affected. Although the phenotype is variable among seedlings from mild to severe, they all exhibit a shorter meristematic zone (Fig.S1 a, b, c). We analyzed in detail the organization of the RAM and root stem cell niche (RSCN) of *drm1;drm2;cmt3* and *nrpd2a;nrpd2b* 10 dpg seedlings and observed that roots from both mutant lines showed a disorganized RAM and defects in the columella region relative to wild-type seedlings (Fig.3). In addition, the loss of function in RBR causes QC divisions, extra stem cells and aberrant divisions and alterations in the columella region, as revealed for the analysis of the *35S::AmiGO-RBR* RAM (Fig.3 e, f). Columella phenotypes observed in RdDM and RBR loss-of-function mutants, shown in Fig.3 and Fig.S1, are consistent with findings in this tissue by Kawakatsu *et al*. (2016), who reported that the *Arabidopsis* columella root cap genome is hypermethylated and transcripts encoding RdDM factors, as well as 24-nt small RNAs (smRNAs), are more abundant in this tissue than any other root cell type.

**Fig. 3.**
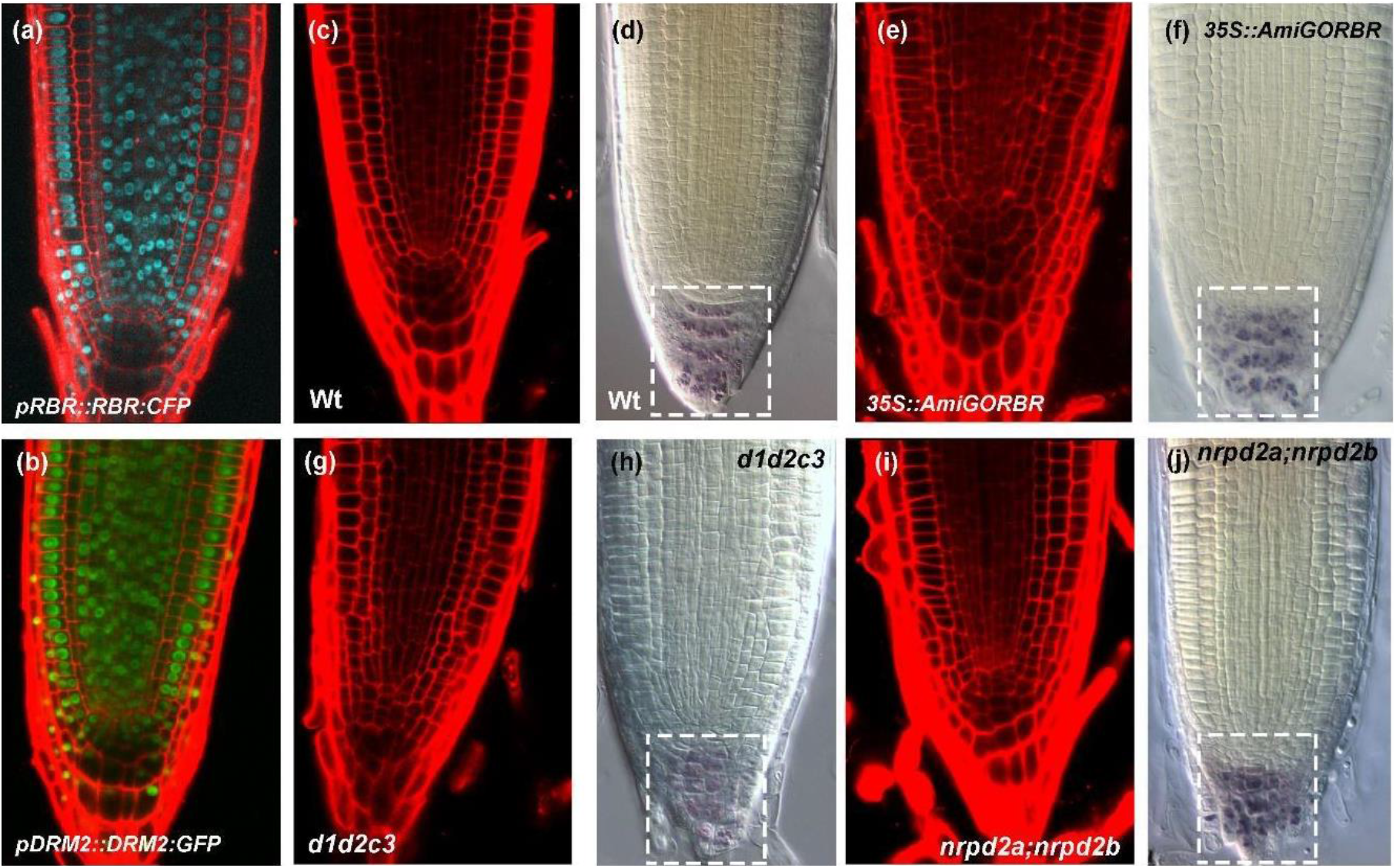
Longitudinal root sections of 10 dpg seedlings imaged by confocal laser scanning microscope (CLSM) (a), (b), (c), (e), (g), (i), and Nomarski optics of lugol-stained roots (d), (f), (h) and (j). Panels (a) and (b) show the expression patterns of *pRBR::RBR:CFP* and *pDRM2::DRM2:GFP*. (c) confocal and (d) Nomarski optics images showing root apical meristem (RAM) and root stem cell niche (RSCN) organization in Col-0 (WT) seedlings. (e) Confocal and (f) Nomarski images of *35S::AmiGO-RBR* seedlings showing alterations in the RAM and RSCN. (g) Confocal and (h) Nomarski images of *drm1;drm2;cmt3* (*d1d2c3*) triple mutant seedlings showing alterations in the RAM and the RSCN. (i) Confocal and (j) Nomarski images of *nrpd2a;nrpd2b* double mutant seedlings showing phenotypes in the RAM and RSCN, dotted squares highlight the Columella region.

### RdDM and SUVR2 targets are up-regulated in the AmiGO-RBR background

SUVR2 silences a subset of RdDM target loci, as well as RdDM-independent targets (Hang *et al*., 2014). Well-known targets of RdDM include TEs from the solo LTR (SLTR) and AtGP1 LTR families and genes such as *SUPPRESSOR OF drm1 drm2 cmt3* (*SDC*) and it has been shown that at *SDC* and *ERT7* loci, the *suvr2* loss-of-function mutants display a synergistic phenotype with mutants in key genes of the RdDM pathway, which suggests that at these loci SUVR2 might exert silencing through a pathway which is partially independent of RdDM (Hang *et al*., 2014). Our data indicates that SUVR2, DRM2 and DRD1 bind *in vitro* to RBR and based on the presence of the LxCxE SLiM other RdDM components like NRPD1, NRPE1 and DCL3 could also potentially bind to RBR. Therefore, we wondered if the silencing of several RdDM and SUVR2 target loci may be affected in a *35S::AmiGO-RBR* background. To answer such question, we isolated total RNA of 12-days-old wild-type and *35S::AmiGO-RBR* seedlings and performed qRT-PCR assays using previously-reported primers for *SDC, AtGP1, solo LTR* (SLTR), AT1TE51360 (AT1TE), AT2TE78930 (AT2TE), ERT7, ERT9, ERT12, and ERT14. Our results showed that all tested loci are either moderately or strongly up-regulated in the RBR loss-of-function background relative to the wild-type control (Fig.4a). It has been shown that *ERT9* transcripts are not de-repressed in the *suvr2* mutant background, suggesting that RBR might influence DRM2 and SUVR2 targets independently. Overall, these results indicate that RBR acts repressing RdDM and SUVR2 transposable elements targets. Whether this action depends on RBR protein-protein interaction with DRM2, SUVR2 or DRD1 remains to be answered in future studies.

**Fig. 4.**
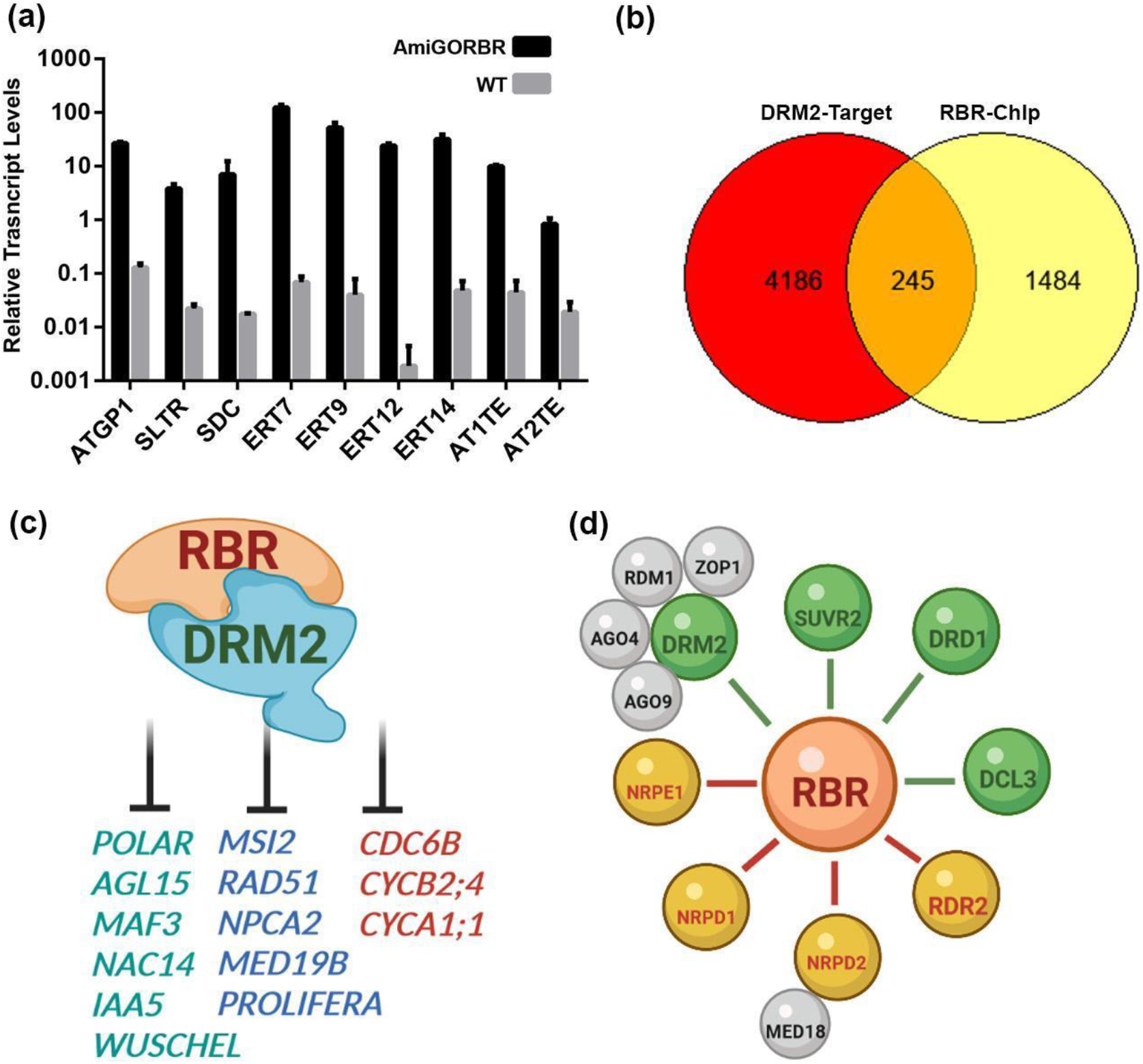
(a) Transcript levels revealed by qRT-PCR of RdDM targets in the AmiGO-RBR mutant background vs the control, (b) RBR-ChIP and DRM2-mediated DNA methylation common targets, (c) Key examples of RBR-DRM2 common targets, (d) RdDM proteins that have been validated as RBR direct interactors (green lines) and those that are putatively binding RBR (red lines), gray balloons are PPIs with DRM2 reported in the IntAct database https://www.ebi.ac.uk/intact/interactions?conversationContext=4, (Created with BioRender.com).

The RdDM pathway methylates not only TE loci, but also hundreds of protein-coding genes (Jha & Shanka, 2014). Since RBR also has hundreds of targets, predicted by Chip-Seq (Bouyer *et al*., 2018), we explored a potential overlap between the 4,431 DRM2 methylation targets proposed by Jha and Shankar (2014) and the 1,729 RBR target genes predicted by Bouyer *et al*., (2018). We found that 245 target genes are shared between RBR and DRM2 (Fig.4b). Among the 245 shared target genes we found several interesting ones (Fig. 4c). We highlighted those that encode transcriptional regulators such as *POLAR LOCALIZATION DURING ASYMMETRIC DIVISION AND REDISTRIBUTION (POLAR), AGAMOUS LIKE 15, NAC15, INDOLE-3-ACETIC ACID INDUCIBLE 5 (IAA5)* and *MADS AFFECTING FLOWERING 3 (MAF3)*. Another important transcription factor that has been shown to act downstream RBR is *WUSCHEL*. In *rbr1-2* mutants, supernumerary megaspore mother cells (MMCs) are formed, a phenotype that correlates with *WUS* transcriptional deregulation Zhao *et al*. (2017). Indeed these authors demonstrate that RBR binds to a specific region on the *WUS* promoter. It has also been shown that in the *drm1;drm2;cmt3 WUS* transcription is de-repressed during root regeneration and that two non-CG sites in the promoter of this gene might be related to *WUS* silencing in *Arabidopsis* roots (Shermer *et al*., 2015). Recently Mendes *et al*. (2020) showed that *drm1;drm2* double mutants develop multiple MMCs, a phenotype also described for other mutants in key genes of the RdDM pathway, such as *rdr6* and *ago9* (Olmedo-Monfil et al., 2010). We also found that genes related to DNA integrity, DNA replication and cell cycle are common targets of RBR and DRM2, such as *RAD51, PROLIFERATING CELL NUCLEAR ANTIGEN 2 (PCNA2), MINICHROMOSOME MAINTENANCE 7/PROLIFERA (MCM7), MS1, MEDIATOR 19B (MED19B), CYCLIN A1;1 (CYCA1;1), CELL DIVISION CONTROL 6B (CDC6B)* and *CYCLIN B2;4 (CYCB2;4)* (Table S2). The putative function of RBR and DRM2 acting on the silencing of genes involved in cell cycle progression, which are normally expressed only in root meristematic cells, such as *CDC6B* or *CYCA1;1*, correlate with some of the root phenotypes reported in Fig. 3 and Fig.S1. However, root phenotypes in RdDM mutants are not similar in all cases, and such contrasting phenotypes may be caused by the deregulation of hundreds of genes with diverse cellular functions.

Overall, this study uncovers novel mechanisms for RBR function in transcriptional silencing through interacting with components of the RdDM pathway and opens novel working hypotheses for diverse potential RBR-RdDM interactions (Fig. 4d), including the RBR-DRM2 complex, regulating TEs and interesting lineage-specific transcription factors.

## Acknowledgements

We wish to thank Vicki Chandler, Steve Jacobsen, Fred Berger and Pauline Jullien for sharing published plant materials. We also thank Dr. Juan Caballero-Pérez for initial advice on bioinformatics. J L-R (CVU 858608) was supported by Consejo Nacional de Ciencia y Tecnología (CONACYT) with a PhD Fellowship. A C-R was supported by EMBO-ALTF 1114-2006 and CONACYT 000000000092916 grants. M A-V was supported by Consejo Nacional de Ciencia y Tecnología (CONACYT) grants 158550 and A1-S-38383 and Newton Fund of the Royal Society grant NA150181.

## Author contributions

Conceived the project: A C-R and M A-V. Performed *wetlab* and *in silico* experiments: A C-R, A E-C, J L-R, I B. Analyzed the data: A C-R, M A-V, B S, I B, A E-C and J L-R. Contributed reagents and equipment: B S, M A-V, A C-R. Wrote the manuscript with inputs from all coauthors: AC-R and MA-V.

## Supporting information

**Fig._S1.**
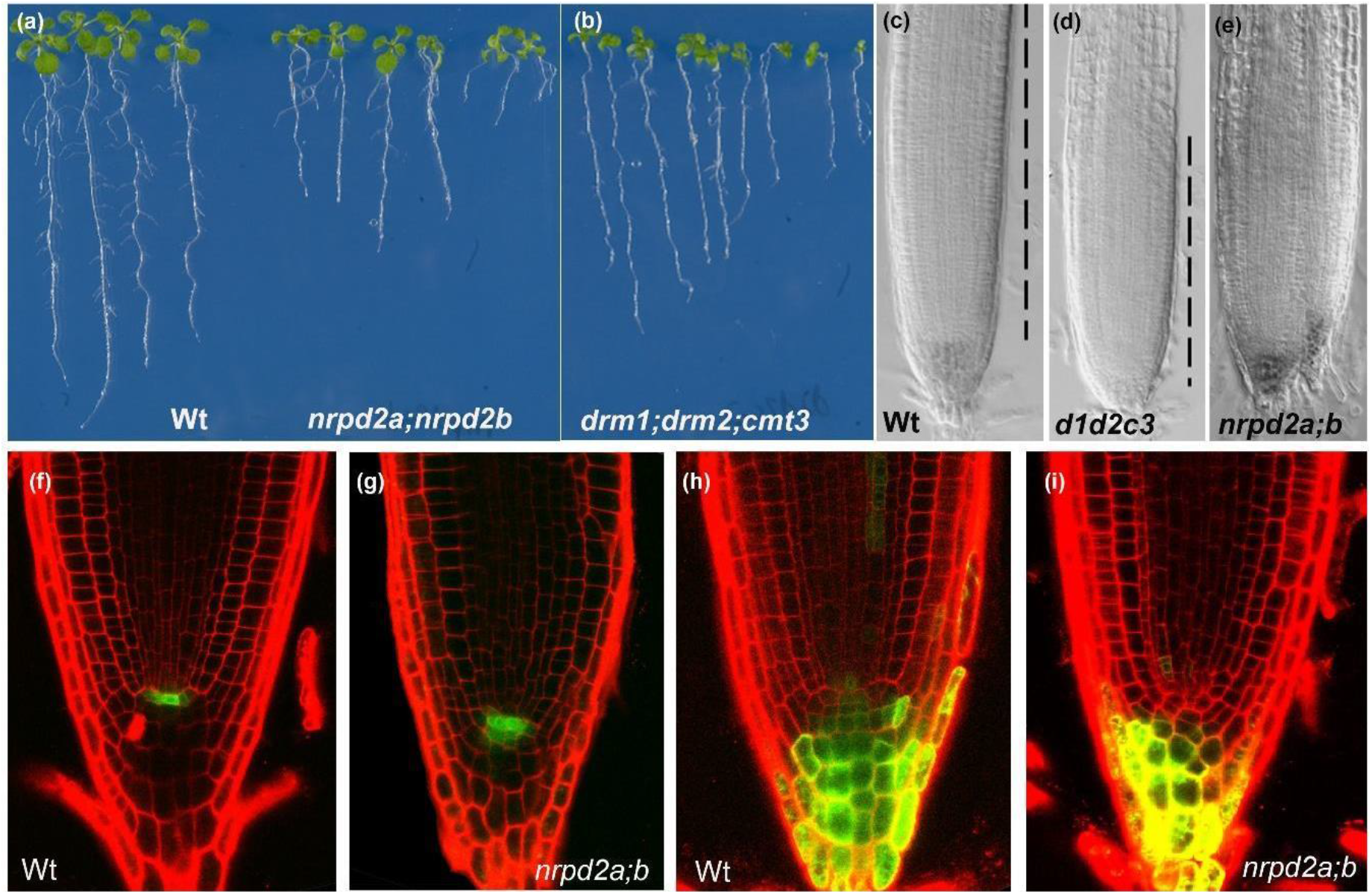
RAM phenotypes in RdDM mutants. Root and shoot phenotypes, recorded with stereomicroscope, of 10 days post germination (dpg) seedlings of wild-type (a), double and triple mutants in RdDM proteins (a), (b). Nomarski optics for RAM phenotypes of *drm1;drm2;cmt3* and *nrpd2a;nrpd2b* mutant (d), (e) and wild-type (c) roots of 10 dpg seedlings. Longitudinal root sections of 10 dpg seedlings by confocal laser scanning microscope (CLSM) of *pWOX5::GFP* and *TCS::GFP* transgenes in WT (f), (h) and *nrpd2a;nrpd2b* (g), (i) backgrounds, respectively.

## Notes

### Competing Interest Statement

The authors have declared no competing interest.

